# Mathematical Modeling of Multiple Pathways in Colorectal Carcinogenesis using Dynamical Systems with Kronecker Structure

**DOI:** 10.1101/2020.08.14.250175

**Authors:** Saskia Haupt, Alexander Zeilmann, Aysel Ahadova, Magnus von Knebel Doeberitz, Matthias Kloor, Vincent Heuveline

## Abstract

Like many other tumors, colorectal cancers develop through multiple pathways containing different driver mutations. This is also true for colorectal carcinogenesis in Lynch syndrome, the most common inherited colorectal cancer syndrome. However, a comprehensive understanding of Lynch syndrome tumor evolution which allows for tailored clinical treatment and even prevention is still lacking.

We suggest a linear autonomous dynamical system modeling the evolution of the different pathways. Starting with the gene mutation graphs of the driver genes, we formulate three key assumptions about how these different mutations might be combined. This approach leads to a dynamical system that is built by the Kronecker sum of the adjacency matrices of the gene mutation graphs. This Kronecker structure makes the dynamical system amenable to a thorough mathematical analysis and medical interpretation, even if the number of incorporated genes or possible mutation states is increased.

For the case that some of the mathematical key assumptions are not satisfied, we explain possible extensions to our model. Additionally, improved bio-medical measurements or novel medical insights can be integrated into the model in a straightforward manner, as all parameters in the model have a biological interpretation. Modifications of the model are able to account for other forms of colorectal carcinogenesis, such as Lynch-like and familial adenomatous polyposis cases.

**Graphical Abstract:**
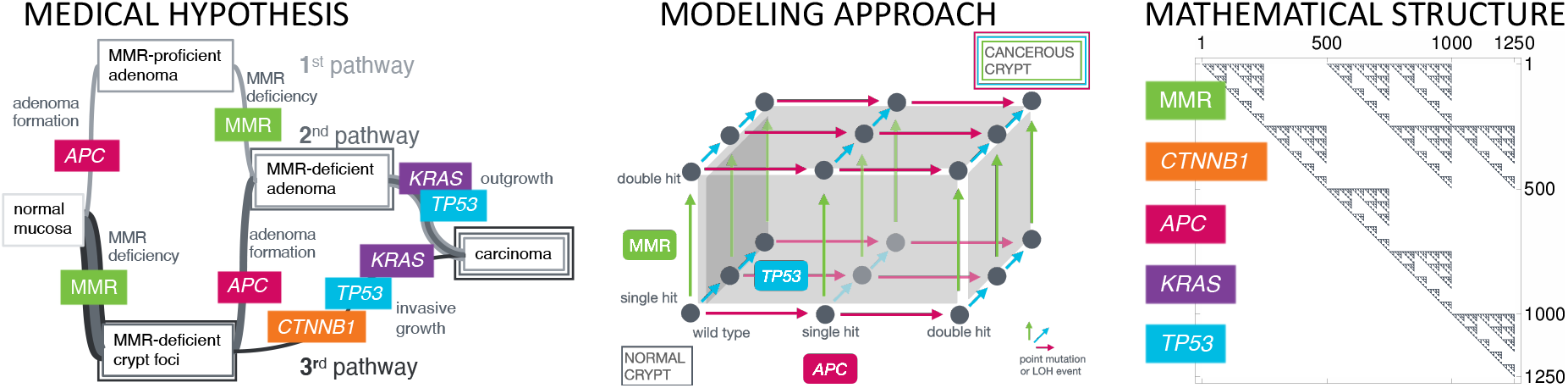
From the Medical Hypothesis Over the Modeling Approach To the Mathematical Structure. The medical hypothesis of multiple pathways in carcinogenesis is widely known for various types of cancer. *Left:* We present a model for this phenomenon at the example of Lynch syndrome, the most common inherited colorectal cancer syndrome, with specific key driver events in the MMR genes, *CTNNB1*, *APC*, *KRAS* and *TP53*. *Middle:* This current medical understanding of carcinogenesis is translated into a mathematical model using a specific dynamical system, which can be represented by a graph structure, where each vertex in the graph represents a genotypic state and the edges correspond to the transition probabilities between those states. Starting with all colonic crypts in the state of all genes being wild-type and a single MMR germline mutation due to Lynch syndrome, we are interested in the distribution of the crypts among the graph at different ages of the patient in order to obtain estimates for the number of crypts in specific states, e.g. adenomatous or cancerous states. *Right:* The underlying matrix of the dynamical system makes use of the Kronecker sum and product. It is a sparse upper triangular matrix accounting for the assumption that mutations cannot be reverted. This allows fast numerical solving by using the matrix exponential. Each nonzero entry of the matrix represents a connection between genotypic states in the graph.

## 1 Introduction

Cancer is the second leading cause of death world-wide accounting for an estimated 9.6 million deaths in 2018, whereby the second most common type is colorectal cancer [Bra+18]. Still, adequate treatment and in particular prevention strategies are lacking in many cases, as it is difficult to measure the process of cancer development, called carcino-genesis, right from the beginning.

In this work, we present a mathematical model of colorectal carcinogenesis. It takes into account the multiple pathway nature of carcinogenesis (Graphical Abstract, left) reflecting different types of colorectal cancer with individual needs for prevention and treatment.

The mathematical model makes use of dynamical systems with a specific matrix structure using Kronecker products and sums (Graphical Abstract, right) in order to systematically describe the mutational events of individual genes (Graphical Abstract, middle).

To account for dependencies between the involved driver mutations, we present several extensions to the model, again by using the Kronecker structure.

To exemplify this approach, we build the model for Lynch syndrome, the most common inherited colorectal cancer syndrome with an estimated population frequency of 1 in 180 [Kli+19]. Lynch syndrome is caused by an inherited mismatch repair (MMR) gene mutation. Colorectal cancers which develop due to Lynch syndrome therefore are MMR-deficient and show increased microsatellite instability (MSI).

In addition to the Lynch syndrome case, we modify the ansatz to model the sporadic counterpart of Lynch syndrome, often called Lynch-like cancers [Car14], as well as the classical adenoma-carcinoma sequence first described by Vogelstein and Kinzler [VK93] for microsatellite-stable (MSS) colorectal cancers. Further, we apply the model to another hereditary colorectal cancer syndrome, familial adenomatous polyposis (FAP) [NA18].

### 1.1 Related Work

First attempts to build mathematical models in cancer research were made in the middle of the 20th century. Armitage and Doll [AD54; AD57] proposed and analyzed one of the first multistage models of carcinogenesis, which are based on the hypothesis that there are multiple subsequent steps before a cancer is formed. The model was extended in the following years [Ken60; Ser84]. Among the first to consider a model of multiple pathways of carcinogenesis were Tan et al. [TB88; TH08]. These are based on the hypothesis that there are several possible ways in which cancer can develop.

With the increasing medical knowledge about cancer development, it became more and more evident that a single model describing the whole process of carcinogenesis from the genomic, over the cell, up to the tissue, organ and population-level is too complex to build. In other words, a single multi-level model of carcinogenesis has not yet been built. Nowadays, there exist different types of models describing individual aspects of carcinogenesis (in an unordered list of example publications):

▷ Modeling **healthy tissue formation**, such as the evolution of colonic crypts [Bin+17; Bak+14; Bak+19],
▷ detecting **driver genes** [Des+99; BES07; Ger+11; Woe+01; Ger+20],
▷ estimating the most likely **temporal order of key mutations** [Tom+15; Mit+18],
▷ modeling the **cancer-immune system interaction**, including neoantigen presentation [BAL18; Lak+19; Bal+19],
▷ predicting **effects of intervention strategies** on tumor growth and patient survival, such as the effect of screening on adenoma risk [Thi+01].

From a mathematical point of view, the modeling makes use of different approaches, such as ordinary differential equations [Kom+02; AGJ08], partial differential equations [Liu+18], stochastic processes [IMN04; Now+02], graph theory [Nax+17; TMS15; Bee+05], and statistics [CZ08; Buc04].

For hereditary colorectal cancers, in particular, Komarova et al. [Kom+02; KSN03] proposed a model for the accumulation and ordering of key events during carcinogenesis based on ordinary differential equations [Kom+02; KSN03], which was adapted to sporadic carcinogenesis. In particular, it addresses the question of to which extent genetic instability is an early event in carcinogenesis.

A recent paper by Paterson et al. [PCB20] is similar in that it solves a stochastic model for quantifying the evolutionary dynamics of colorectal cancer initiation and progression.

### 1.2 Contribution

Our model is similar to the latter two in the sense that it aims to describe the accumulation of key drivers in colorectal carcinogenesis by using ordinary differential equations. Instead of focusing on modeling *APC* inactivation and mismatch repair deficiency as in [Kom+02], we choose a more general approach for combining mutations in different genes. Compared to [PCB20], we take into account different modes of cancer evolution beside the classical adenoma-carcinoma sequence of colorectal carcinogenesis, including hereditary forms like Lynch syndrome and familial adenomatous polyposis.

We formulate the medical assumptions mathematically leading to the Kronecker structure used for the system matrix in the dynamical system. The use of the Kronecker structure in our model is essential for a systematic analysis of the incorporated genes and bio-molecular effects. It not only makes the model more interpretable from a medical point of view but also reduces the computational costs tremendously. Further, all parameters in the model have a medical interpretation which eases the validation with available data.

Our approach makes it possible to easily extend the model with new medical insights, while preserving the other properties of the model, like the integration of the involved differential equations.

Finally, we solve the ordinary differential equations analytically by using the matrix exponential instead of considering approximations to the matrix exponential series or using stochastic simulations.

### 1.3 Organization

To make this paper self-contained, we elucidate the medical and mathematical background in Sections 2 and 3. The modeling starts in Section 4, where we present our basic model for colorectal Lynch syndrome carcinogenesis. This basic model serves as the groundwork for a variety of extensions which are discussed and analyzed in Section 5. Section 6 represents modifications for non-Lynch scenarios or cancers in other organs than the colon. Section 7 demonstrates a selection of the results which can be obtained with the basic model, its extensions and modifications. Finally, we conclude in Section 8.

## 2 Medical Background

Cancer is a disease caused by alterations of the genome, the carrier of genetic information [Wun06; EK05]. Precisely defining these changes, which are required to transform a normal cell of the human body into a malignant tumor cell, is a crucial step towards understanding the development of cancer.

### 2.1 Multiple Pathways in Carcinogenesis

In the early stages of cancer research, it was unknown whether the development of cancer, a process called carcinogenesis, was a purely chaotic process of random mutations. However, in 1959, Nowell and Hungerford [NH04] made the observation of a specific recurrent alteration across different cancers of the same type. This suggested the existence of at least a certain degree of order in the chaos.

In the following decade, evidence emerged that one single mutation is normally insufficient to drive a cell into malignancy because cells possess multiple control mechanisms which protect the organism from the uncontrolled growth of single cells. Thus, Vogelstein, Fearon and Kinzler [Vog+88; VK93] established a step-wise hypothesis of cancer formation postulating that several mutations are required for the development of tumor cells. This hypothesis has been validated subsequently in many independent studies for many different cancer types. Currently, it is expected that a minimum number of three mutation events is required to transform a normal cell into a cancer cell. This hypothesis is called the three strikes hypothesis [Tom+15].

Mutations occur over the whole genome, whereby we differentiate between two broad classes: So-called point mutations only affect a single nucleobase, while loss of heterozygosity (LOH) refers to the loss of larger regions of a chromosome, which can result in the deletion of whole genes.

As about 99 % of the genome is not translated into proteins, most mutations do not have a direct consequence on cell viability or behavior. However, if mutations strike in regions with a protein-encoding function, two main scenarios that can favor uncontrolled cell growth are seen: Mutations can either activate oncogenes (called gain-of-function mutations), which normally promote appropriate cell growth, or mutations can damage or destroy tumor suppressor genes (called loss-of-function mutations), which normally limit cell growth.

This means that, from all the possible mutations that can occur, we need to identify the relatively few key events which have a functional impact on the cell. This includes the identification of oncogenes and tumor suppressor genes, but there are many more mutations to be identified. Moreover, only a certain combination of these mutations will lead to cancer in the end. This is due to the fact that some mutations have a growth-repressing effect and lead to cell death.

Different combinations of key mutations result in several distinct pathways. An important goal in cancer research is to investigate which of these pathways can arise in human carcinogenesis. This leads to a high-dimensional and complex problem which requires a mathematical description of carcinogenesis including all the mutational events. In this paper, we propose a novel approach to tackle this problem which is strongly driven by its medical interpretability.

### 2.2 Lynch Syndrome-Associated Colorectal Carcinogenesis

Central to carcinogenesis is the accumulation of key mutations in driver genes. Mutations are errors which occur during DNA replication within cell division and which are not repaired by one of the error detection, repair and control systems present in all organisms. Central to this essential network, which enables healthy life, are DNA repair enzymes. Even when repair enzymes work properly, some alterations will always be detectable in newly generated cells. If DNA repair itself is impaired, the number of these mutations will rise dramatically.

This is particularly true for patients with a deficient DNA repair mechanism, being unable to repair errors which occur during DNA replication [Kol96]. This arises when patients have mutations on both alleles in one of the four underlying mismatch repair genes *MLH1*, *MSH2*, *MSH6* and *PMS2* [Cha03] leading to MMR deficiency. Then, DNA replication errors, especially those which occur at repetitive sequences (the same base pairs occur successively, called microsatellites) cannot be corrected by the mismatch repair system. In other words, MMR deficiency leads to microsatellite instability.

Most of the cancers in the general population occur by chance. These cancers are called sporadic. However, in some families, certain types of cancer appear more frequently. This is either a familial or a hereditary form of cancer. The former is due to a combination of genetic and environmental factors but in contrast to hereditary cancers, there is not a specific pattern of altered genes which is passed down in the family from parent to child. From a modeling point of view, the advantage of focusing on hereditary tumors is that there are clearly defined molecular events determining the onset of the disease and thereby representing a known mechanism underlying carcinogenesis.

In this work, we focus on modeling the carcinogenesis in Lynch syndrome, which is the most common inherited colorectal cancer syndrome [Cha03].

Lynch syndrome carriers are predisposed to develop MSI cancers due to an inherited pathogenic variant in one allele of the affected MMR genes. Upon the second somatic hit inactivating the remaining allele, MMR deficiency manifests in the affected cell.

Individuals with Lynch syndrome are predisposed to developing certain malignancies with a substantially higher lifetime risk compared to the general population. The most common Lynch syndrome manifestations are colorectal cancer (CRC, 60–80 % [KKG05] compared to 6 % in the normal population) and endometrial cancer (50–60 % compared to 2.6 % in women without Lynch syndrome) [Jas+10; Kaa+16]. Further, individuals have an increased lifetime risk for many other types of cancer such as in the stomach, small bowel, brain, skin, pancreas, biliary tract, ovary (only for women) and upper urinary tract [KKG05].

Among other cell structures, the human colon consists of colonic crypts which are found in the epithelia of the colon and which consist of different cell types [Coo19]. If a cell in a crypt becomes mutated, this mutation has to spread within the crypt such that the whole crypt is mutated and can be measured with current techniques. In order to compare our modeling results with currently available data, we focus on the evolution of genetic states within crypts as a whole.

For colorectal cancer in general, there is one dominant hypothesis for carcinogenesis, which was first proposed in the late 1980s by Vogelstein, Kinzler and Fearon [Vog+88; VK93]. This Adenoma-Carcinoma Hypothesis describes the formation of certain precancerous lesions and their progression into a manifest cancer. The model implies that adenomas are the precursor lesions of most colorectal cancers and it describes typical molecular events associated with tumor progression.

In addition, in Lynch syndrome patients, cancers occur [Eng+18] without an adenoma phase which leads to an adaptation of the model with different pathways of Lynch syndrome colorectal carcinogenesis [Aha+18]. As explained in Section 2.1, key events in carcinogenesis are the mutations in tumor suppressor genes and oncogenes. In Lynch syndrome, an additional key event is MMR inactivation, which occurs due to a second somatic mutation in one of the four MMR genes. Altogether, this leads to the current Three Pathway Hypothesis of Lynch syndrome cancer development formulated in [Aha+16; Aha+18] and illustrated in the graphical abstract. The relative proportion of one or the other pathway of progression and the contribution of certain molecular events is thereby an open question to be addressed by mathematical cancer modeling. By proposing a mathematical modeling ansatz for describing the multiple pathways in Lynch syndrome carcinogenesis, we investigate the distribution among the different genotypic states and pathways in order to address these open questions.

## 3 Mathematical Background

In this section, we will give a short introduction to the mathematical background of our model. We start with a few basic notions from graph theory, before we state three assumptions about the properties a combination of several processes should have. The assumptions are then related to graph products and the Kronecker sum of matrices. Finally, we consider linear dynamical systems and their solution.

### 3.1 A Primer on Graph Theory

Modeling evolutionary processes can often be done with the help of graphs 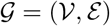, i.e. as set 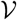 of states (vertices) which are combined with edges *ε* if a progression between the states is possible.

We represent the graph as an adjacency matrix *A*, that is, a matrix which has as many rows and columns as there are vertices in the graph. The entry *a*_*i,j*_ at position (*i, j*) of the matrix *A* indicates whether there is no connection (*a*_*i,j*_ = 0), a weak relationship (*a*_*i,j*_ is small) or a strong bond (*a*_*i,j*_ is large) between the vertices *i* and *j*. The value *a*_*i,j*_ is often referred to as the weight of the edge *ij*. Vertices *i* and *j* which are connected by an edge are called adjacent and are denoted by *i* ~ *j*. Vertices may also be connected to themselves. In this case, the edge is called a self-loop.

It is also possible to indicate directed edges, by letting *a*_*i,j*_ (weight of the edge from *i* to *j*) differ from *a*_*j,i*_ (weight of the edge from *j* to *i*). A directed graph with no directed cycles is called a directed acyclic graph (DAG). The vertices of a DAG can be ordered such that the adjacency matrix is an upper triangular matrix.

### 3.2 Combining Processes and the Cartesian Product of Graphs

Now assume that we have two processes, e.g. the accumulation of mutations in two different genes of the same cell, which we want to combine and represent as one single process. For this combination we have the following key assumptions:

**Existence of States** The states in the combined process should exactly be all possible combinations of states from the underlying processes.
**Edge Connectivity** We require that in the combined model only one of the processes can evolve to the next state at any point in time.
**Independence of the Processes** We require that the processes are independent of each other.

In the context of the example above, we interpret from the first assumption that all combinations of mutations in the different genes are possible (i.e. there are no mutations that prevent other mutations) and that there are no additional states. This also implies that the order in which mutations are accumulated is ignored.

The edge connectivity assumption states that no two mutations in different genes can occur at the exactly same point in time.

The third assumption entails that a mutation in one gene does not change the mutation probability in another gene. While the independence is a strong requirement, it makes the mathematical analysis more amenable by reducing the number of parameters and improving the model structure. However, even in the case where the independence requirement is not justified or its justification is unknown, it can serve as a valuable baseline when figuring out in which parts of the model and to which degree the independence is violated. We follow this chain of thought further in the extensions of our basic model in Section 5.

Consider two processes that are represented by the graphs 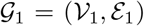 and 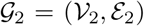. The combined process, which satisfies our key assumptions, is then represented by a 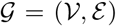, that is given by the Cartesian product [HIK11] of the graphs 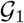 and 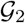

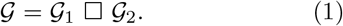

The set of vertices 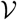 of the Cartesian graph product is given by the Cartesian product 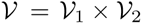 of the vertex sets 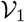 and 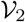. This means the vertex set 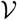 consists of all possible combinations of vertices from the first graph with vertices from the second graph

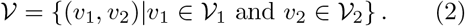

In other words, the first requirement is satisfied.

The edge set *ε* is made up of edges of the forms

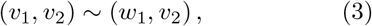

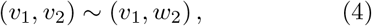

where the vertices 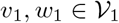 are adjacent in the graph 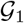 and similarly for 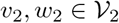. This means that we connect the states in our combined process such that each edge corresponds to a single edge in exactly one of the underlying processes. From this we establish that our second assumption is also fulfilled.

As each edge in *ε* corresponds to exactly one of the edges in *ε*_1_ ⋃ *ε*_2_, we can transfer the edge weights from the graphs 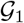 and 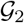 to 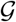. From this, we conclude the satisfaction of the independence assumption.

We can extend this combination to more than two processes by iteratively applying the definition to two of the processes.

To relate the Cartesian product of graphs to their adjacency matrices, we first need to consider the Kronecker product and sum of two matrices.

### 3.3 Kronecker Product and Sum

The Kronecker product 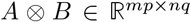 of two matrices 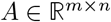 and 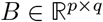 is defined [HJ91; Loa00] by the block matrix

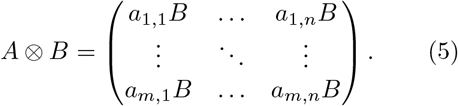

For square matrices 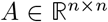 and 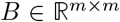, we further define the Kronecker sum

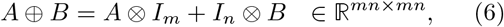

where *I*_*m*_ (resp. *I*_*n*_) denotes the identity matrix of size *m* × *m* (resp. *n* × *n*).

The Kronecker product and sum possess several favorable properties, among which is the compatibility with the matrix transpose

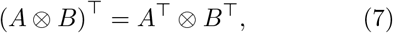

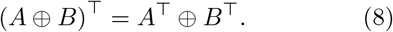

Let *A*_1_ and *A*_2_ be the adjacency matrices of the graphs 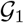 and 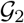. The adjacency matrix of the Cartesian Product 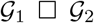 is given by the Kronecker sum *A*_1_ ⊕ *A*_2_ [KR05].

This connection between the Cartesian product of graphs and the Kronecker sum is visualized in Figure 1.

**Figure 1:**
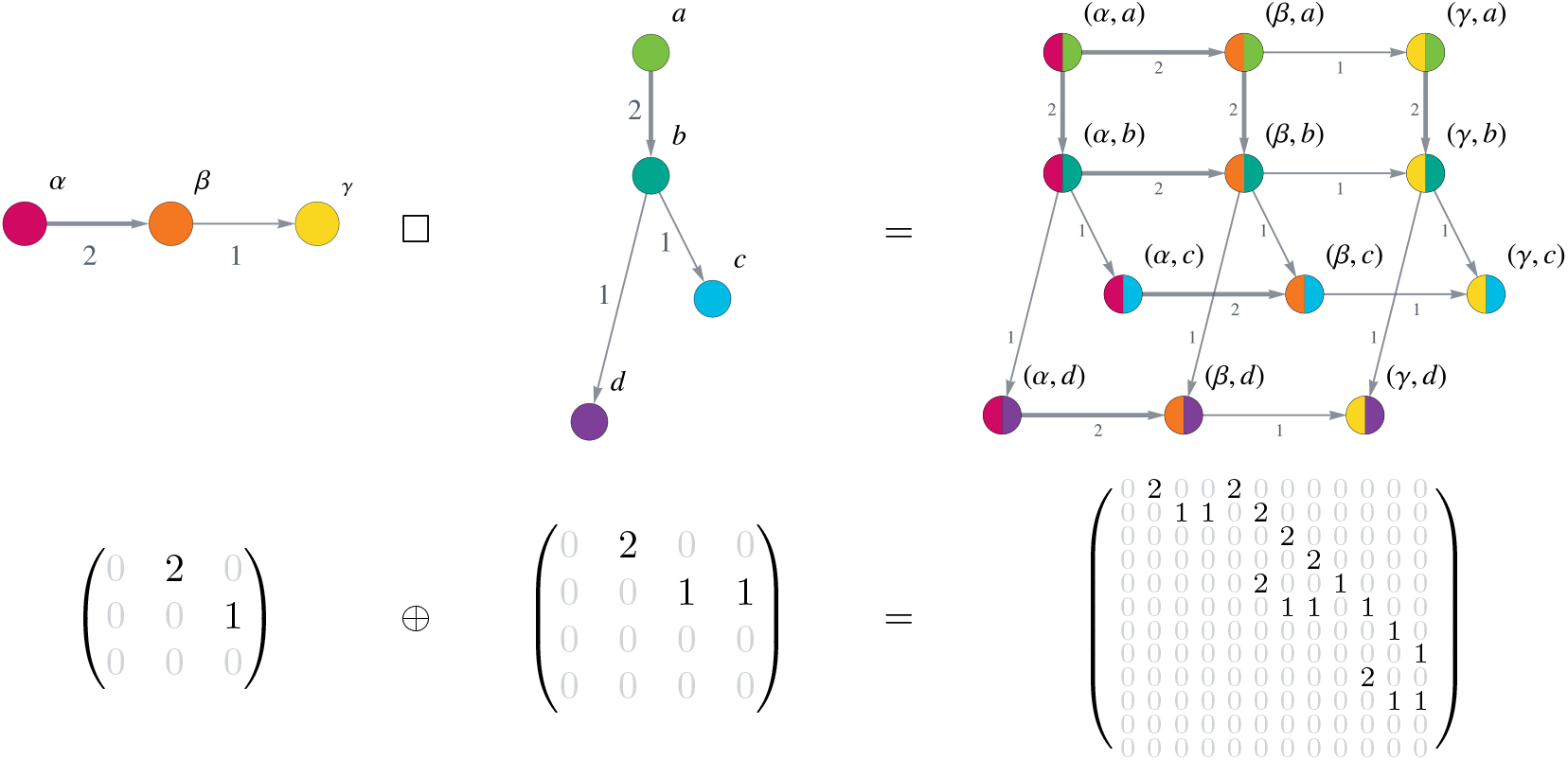
Cartesian Product of Graphs. The upper row shows two graphs and their Cartesian product. Notice how each vertex (*α, β, γ*) of the first graph is combined with each vertex (*a, b, c, d*) from the second graph, yielding a total of 12 (= 3 · 4) vertices in the Cartesian graph product. The edge weights (indicated by numbers next to the edges and the edge thickness) of the graphs on the left and middle transfer to the corresponding edges in the product graph. The bottom row displays the adjacency matrices corresponding to the graphs in the upper row as an equation involving the Kronecker sum of the matrices.

The Kronecker product is not commutative, i.e. in general we have *A*_1_ ⊕ *A*_2_ ≠ *A*_2_ ⊕ *A*_1_. Accordingly, the graph products 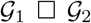 and 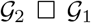 are not equal to each other. However, as they are isomorphic to each other, we are free to choose an ordering of the matrices as long as we use the same ordering in all computations.

### 3.4 Linear Dynamical Systems

An important and highly used class of equations for dynamical systems in mathematical modeling are linear differential equations of the form

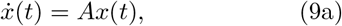

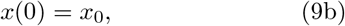

where 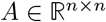 is a matrix (later chosen to be the adjacency matrix of a graph), which is constant with respect to time, and 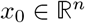 is a vector representing the initial value.

Linear differential equations have a unique solution [Tes12, p. 60], which is given by

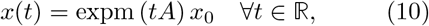

where expm(*A*) describes the matrix exponential of the matrix *A*, which is defined by

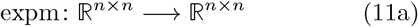

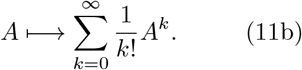

Computing the matrix exponential [MV03] can be done in a variety of different ways. However, as the matrix exponential is multiplied with the vector *x*_0_, we do not need to compute the matrix exponential as a full matrix, but only the action of the matrix exponential on the vector *x*_0_. Algorithms for this task are studied in the context of exponential integrators [AH11; NW12].

The definition of the matrix exponential directly yields that the matrix exponential of a triangular matrix (as it is in our case) is again a triangular matrix.

Later in our basic model (see Section 4.5), the matrix *A* in the linear dynamical system will be the Kronecker sum of several matrices (27). The matrix exponential of such a matrix simplifies according to [Hig08, Theorem 10.9, p. 237] to

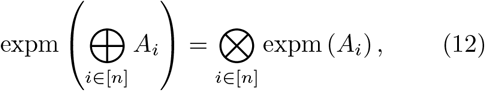

where [*n*] denotes the set of integers from 1 to *n*.

Thus, instead of computing the matrix exponential of one large matrix, we can compute the matrix exponentials of several small matrices and connect them with the Kronecker product, which gives an additional performance boost for numerical computations.

The equation (12) can be seen as a generalization of *e*^*a*+*b*^ = *e*^*a*^*e*^*b*^ for real numbers *a* and *b*. However, it is important to note that this statement does not hold for the standard matrix addition and product. Thus, in general, we have [Hig08, Theorem 10.2, p. 235]

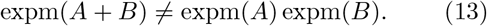

Later, in the extensions of our model in Section 5, we consider sums of matrices (32).

While in this case the matrices cannot be simplified as much as for our basic model, we can still use the exponential integration mentioned above. Further mathematical analysis is however possible in the context of operator splitting [MQ02] or the Baker-Campbell-Hausdorff formula [BB18].

Often, the initial value *x*_0_ itself can be written as a Kronecker product, i.e.

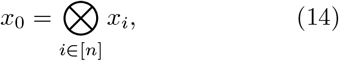

where the *x*_*i*_ are vectors with sizes corresponding to the sizes of the matrices *A*^*i*^. With this, the solution of the dynamical system (9) can be written as

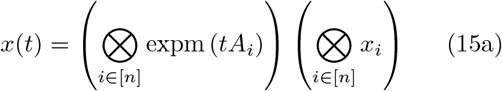

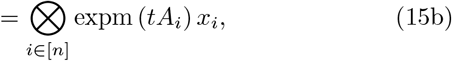

where the last equality is due to the mixed product property of Kronecker products [Loa00].

In most cases, we are not only interested in a single entry of the solution vector *x*(*t*), but in the sum of several entries. To achieve this, we consider the scalar product *v*^⊤^*x*(*t*) of the vector *x*(*t*) with a vector *v* of the same size which has a 1 in all states we want to consider and a 0 everywhere else.

Often this vector can, similarly to the initial value *x*_0_ in equation (14), be written as the Kronecker product of vectors *v*_*i*_ with sizes corresponding to the matrices *A*_*i*_

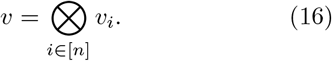

In this case, the accumulation simplifies to

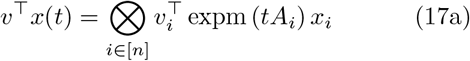

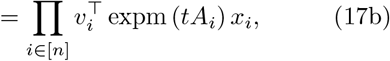

where the first equality follows as above from the mixed product property of Kronecker products and the second one is due to the fact that the Kronecker product of real numbers (here: 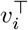 expm (*tA*_*i*_) *x*_*i*_) is the standard product of real numbers.

## 4 Modeling Lynch Syndrome Carcinogenesis

In this section, we introduce our basic model for modeling colorectal carcinogenesis in Lynch syndrome. The model consists of a dynamical system given in the form of a linear ordinary differential equation which is constructed with the help of an adjacency matrix describing the joint process of mutations in several genes. Those mutations are assumed to be present in the whole crypt. Mutations which occur in one cell but are washed out as they reach the top of the crypt and undergo apoptosis are not considered in the model.

### 4.1 Gene Mutation Graphs

In the case of colorectal carcinogenesis in Lynch syndrome, the MMR gene mutations are modeled as the causative mechanism for the increased cancer lifetime risk of Lynch syndrome individuals. Besides the MMR genes, we consider four additional possible driver genes, namely *APC*, *KRAS*, *CTNNB1* and *TP53* which are typical representatives of the oncogenes and tumor suppressor genes affected in the corresponding pathways.

Each of these genes can be in a variety of mutation states:

**State ∅:** In this state, none of the alleles has a point mutation or is affected by an LOH event.
**States `m` and `mm`:** These states describe one allele being hit by a point mutation (where the other one is not mutated) and point mutations on both alleles.
**States `l` and `ll`:** Similarly, these states describe one (respectively two) alleles being affected by an LOH event.
**State `ml`:** One of the alleles has obtained a point mutation and in the other one, an LOH event occurred. We do not differentiate which allele has which mutation and in which order they happened.

We assume that `ll` in *CTNNB1*, *APC* and *TP53* damage a cell in such a way that it directly leads to cell death. Thus, there will be no crypt with all cells being in that state. As we model the evolution of genotypic states of crypts, we do not consider the `ll` states for *CTNNB1*, *APC* and *TP53*.

As we model Lynch syndrome carcinogenesis, all cells and hence, also all crypts have a single germline mutation in the respective MMR gene and there is no ∅ state for MMR.

Further, *APC* and *TP53* are tumor suppressor genes meaning that both alleles have to be mutated for an inactivation. On the other hand, *KRAS* is an oncogene, where one activating mutation is necessary. In colorectal cancer, biallelic mutations of *CTNNB1* seem to be required to mediate an oncogenic driver effect [Hue+15].

All these assumptions lead to the vertex sets

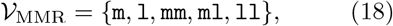

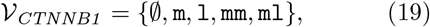

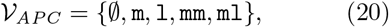

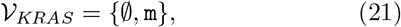

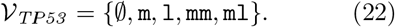

Using these vertex sets, we construct gene mutation graphs, in which we connect the mutation states that differ by only one mutation. This means we assume that only one mutation happens at any specific time point.

Further, we make the assumption that once a mutation has happened it cannot be undone by another mutation. Because of this, the mutation graphs are directed acyclic graphs and their adjacency matrix can be written as a triangular matrix.

The resulting graphs are illustrated in Figure 2. This figure also displays the edge weights of the gene mutation graph, i.e. the likelihood that we transfer from one mutation state to another. The choice of the edge weights will be explained in the following sections.

**Figure 2:**
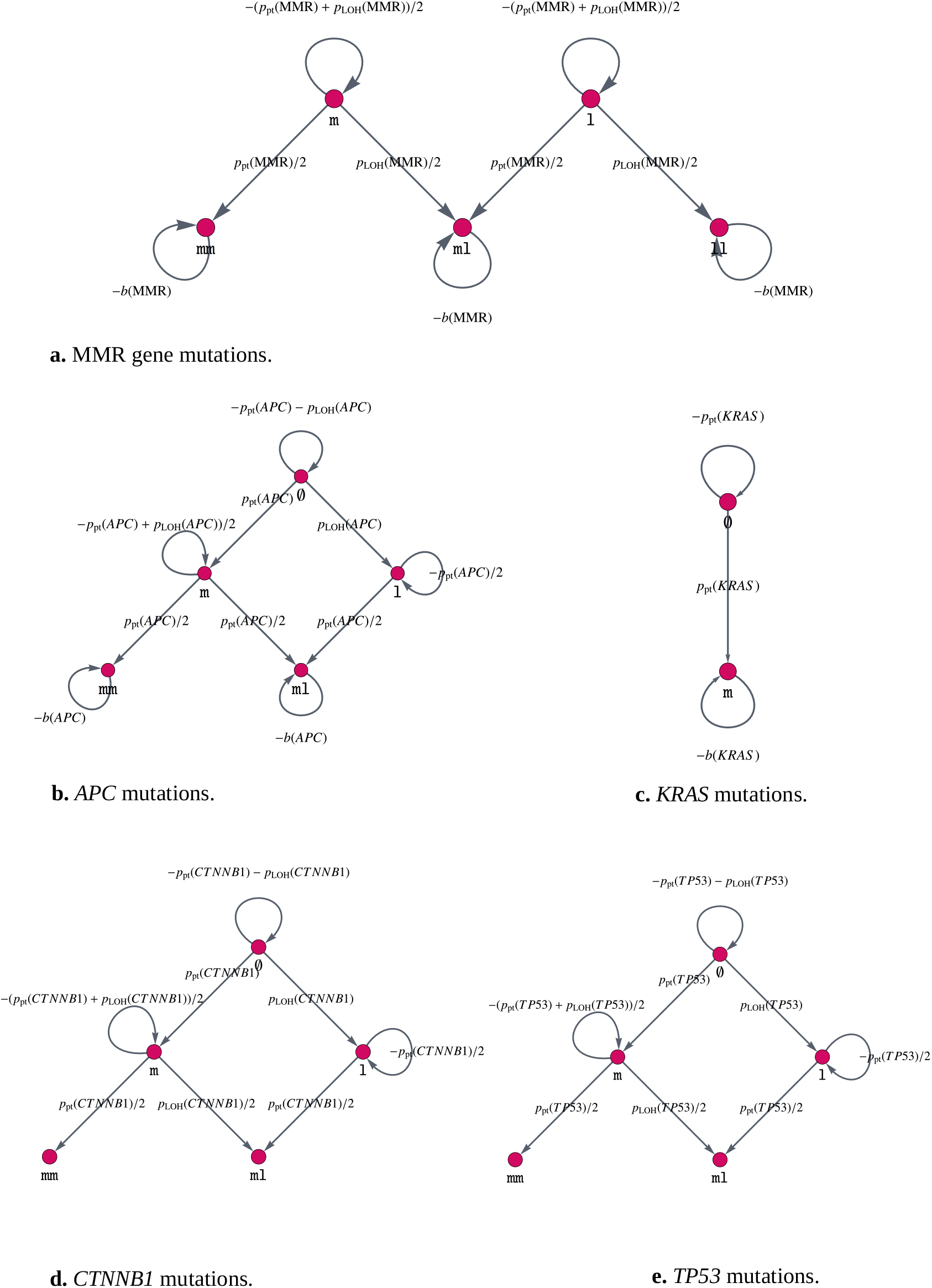
Gene Mutation Graphs. These graphs represent the possible mutation states, i.e. which mutations the alleles of the gene can have accumulated, as vertices ∅, `m`, `l`, `mm`, `ll` and `ml`. The edges connecting different vertices represent mutations, whereas self-loops, i.e. edges that connect a vertex with itself, describe no mutation occurring at the current point in time. The edges are labeled by the amount of change which happens at each point in time.

### 4.2 Point Mutations

To model the likelihood *p*_pt_(gene) for crypts being affected by point mutations in a specific gene, we make the following configurable assumptions:

▷ We would like to model the evolution of crypts over years. Many measurements and estimates are given in days. Thus, we use the factor 365 to convert the measurements per day to measurements per year.
▷ In each cell division, we accumulate *n*_pt_ = 1.2 point mutations according to measurements in [Wer+20], where we assume that a cell division takes one day [Coo19].
▷ The point mutations are uniformly distributed over the base pairs on the entire genome.
▷ Each crypt is estimated [Bak+14] to consist of approximately 1.7 · 10^3^ to 2.5 · 10^3^ cells, whereas only approximately 75 % of them can divide. Thus, we use *n*_cells_ = 1500 as an approximation to the number of cells per crypt.
▷ There are *n*_bp, genome_ = 3.2 · 10^9^ base pairs (bp) on the genome.
▷ Only the point mutations which occur in hotspots of the genes are relevant for the development of a tumor. Hotspots are regions of a gene which give rise to a phenotypical change if mutated. The size of the hotspots *n*_hs_(gene) is gene dependent and is explained in the following.
▷ Not all point mutations which appear in a crypt take over the entire crypt [Nic+18]. We model this in a gene dependent fixation affinity *f* (gene), i.e. the tendency of a cell with a mutation in a gene to take over the whole crypt.
▷ We assume that the alleles are independent of each other, i.e. a mutation in one allele does not influence the mutation probability in the other allele. Thus, the likelihood *p*_pt_(gene) is twice as large if there is no mutated allele (*n*_mut_(gene) = 0) compared to the state where one allele is already mutated (*n*_mut_(gene) = 1).

These assumptions lead to the following formula for the likelihood *p*_pt_(gene):

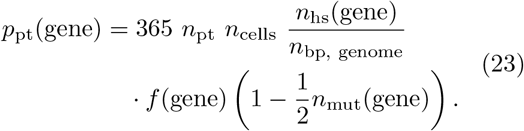

Regarding the hotspots, we assume for *MLH1* and *MSH2* that the whole coding sequence is susceptible to inactivating point mutations, where we use the reference sequence database at NCBI for coding sequence lengths [OLe+16]. For *APC*, we use mutation data from the publicly available DFCI database using the cBioPortal website [Cer+12; Gao+13].

We make use of data from about 4000 CRC samples to identify approximately 2400 hotspots.

For the present model, we assume for *CTNNB1* that only 4 mutations in codon 45 are relevant, according to [Kus+02]. In summary, we obtain the following numbers for *n*_hs_ given in Table 1.

**Table 1:** The following **estimates for** *n*_**hs**_ are used for the computation of the point mutation rates for the individual genes.

| gene | $n_{\text{hs}}(\text{gene})$ |
| --- | --- |
| *MLH1* | 2,270 |
| *MSH2* | 2,800 |
| *CTNNB1* | 1 |
| *APC* | 2,400 |
| *KRAS* | 7 |
| *TP53* | 1,180 |

### 4.3 LOH Events

We assume that all detectable LOH events are large enough to inactivate an affected gene. In other words, we assume that if LOH affects a certain gene, then an exon will be lost and the gene, therefore, is inactivated. As a consequence, the probability of LOH *p*_LOH_(gene) for a given gene is proportional to its length, denoted by *n*_bp_(gene).

The probability of a relevant LOH event for a specific gene with *n*_mut_(gene) ∈ {0, 1, 2} already mutated alleles and length *n*_bp_(gene) bp to be present in the whole crypt is given by

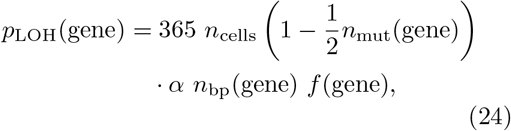

where 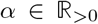 is a parameter to be estimated, independent of the considered gene.

The available data for *MLH1* suggests that inactivation is twice as likely to occur due to LOH than due to point mutations [Por+17]. Thus, we assume

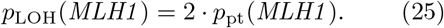

Together with (23) and (24), we get

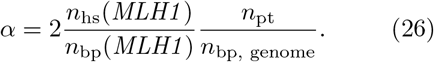

In order to determine *α* and *p*_LOH_, we again use the reference sequence database at NCBI for the length of individual genes [OLe+16] given in Table 2.

**Table 2:**
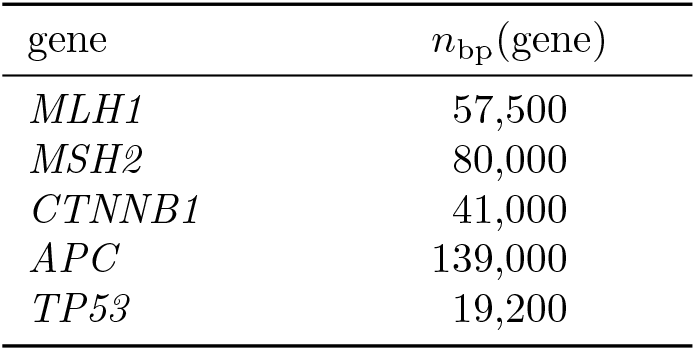
The following **estimates for** *n*_**bp**_ are necessary for the computation of the LOH rates for the individual genes.

| gene | $n_{\text{bp}}(\text{gene})$ |
| --- | --- |
| *MLH1* | 57,500 |
| *MSH2* | 80,000 |
| *CTNNB1* | 41,000 |
| *APC* | 139,000 |
| *TP53* | 19,200 |

### 4.4 Fitness Advantages and Clonal Expansion

There is the possibility of introducing fitness changes *b*(gene) for individual mutation states of a gene. As we model the evolution of mutations at the crypt level, this corresponds to the clonal expansion of the crypts with one of the considered mutations. A fitness advantage is ensured by *b*(gene) > 0 and a disadvantage with *b*(gene) < 0. By using the notion of graphs, this corresponds to a self-loop of the respective genotypic state node with a weight equal to the fitness change. We assume that MMR deficiency leads to a fitness disadvantage [GGS20], i.e. *b*(MMR) < 0, and *APC* inactivation and *KRAS* activation lead to a fitness advantage, i.e. *b*(*APC*) > 0 and *b*(*KRAS*) > 0, in concordance with current measurements [BG16; Nic+18].

### 4.5 A Basic Model for Carcinogenesis

Having defined the gene mutation graphs with adjacency matrices *A*_MMR_, *A*_*APC*_, *A*_*KRAS*_, *A*_*CTNNB1*_, *A*_*TP53*_ for different genes, we combine them as explained in Section 3.2. Accordingly, the adjacency matrix of the combined model is given by the Kronecker sum of the adjacency matrices of the individual genes

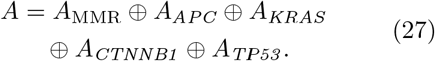

While the matrix *A* has 1250 = 5 · 5 · 2 · 5 · 5 rows and columns, corresponding to all possible genotypes, it is very sparse, which is illustrated in Figure 3(a).

**Figure 3:**
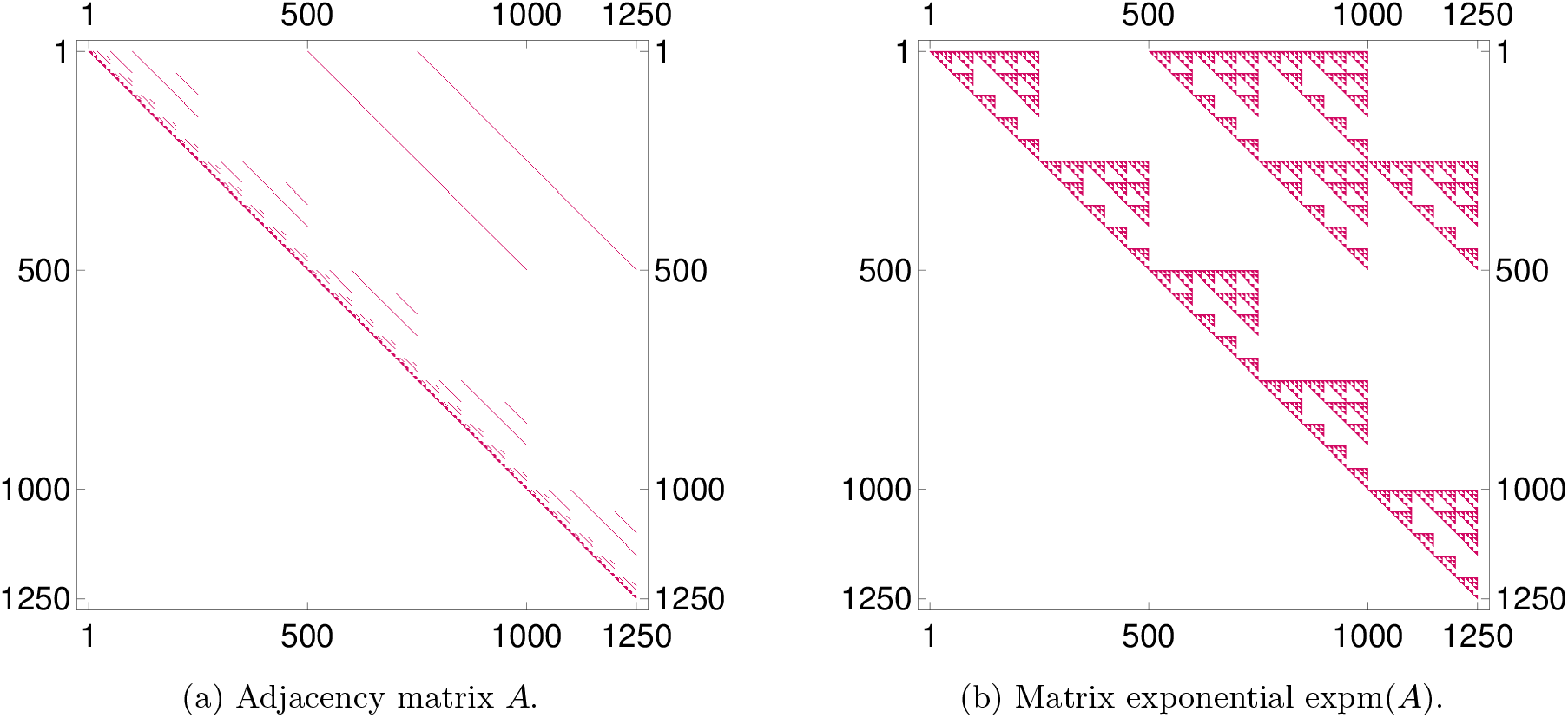
Sparse Matrix Structure. The adjacency matrix *A* (on the left) of the basic model is a very sparse matrix, i.e. only a few entries are nonzero. These nonzero entries are colored red in the plot, which also illustrates the fact that *A* is an upper triangular matrix. The sparsity structure (on the right) of the matrix expm(*A*), which is reminiscent of a Sierpiński fractal, is due to *A* being the Kronecker sum of matrices. The two plots also illustrate nicely how modeling sparse local interactions in the matrix *A* can have a more global effect in expm(*A*).

We use the transpose of this matrix as a system matrix to construct a linear differential equation

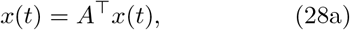

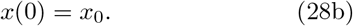

The transpose of the matrix is merely due to different notation conventions for adjacency matrices and differential equations.

We assume that the Lynch syndrome patients have no mutations at birth except for an MMR germline mutation due to a point mutation (90– 95 % of patients) or due to an LOH event (5–10 % of patients) [Klo+12]. We differentiate these two groups of patients by using different initial values for the differential equation. The initial value *x*_0_ for the first group of patients is

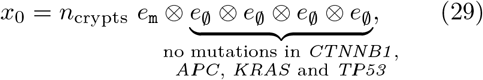

where *n*_crypts_ = 9.95 · 10^6^ is the estimated [HD02] number of crypts in the colon and *e*_m_ denotes the unit vector, which is zero everywhere, except for a 1 at the entry corresponding to the mutation `m`. This initial value can also be described as a vector which has the entry *n*_crypts_ at the position corresponding to the genotype (`m`, ∅, ∅, ∅,) and is zero everywhere else.

Accordingly, the initial value for the second group of patients is given by

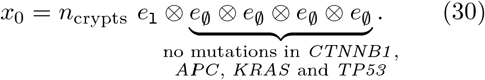

As stated in (10), the exact solution of the differential equation is given by *x*(*t*) = expm(*tA*^⊤^)*x*_0_. We illustrate the sparsity structure of the matrix exponential in Figure 3(b).

According to equation (15), the solution can be rewritten in the following way

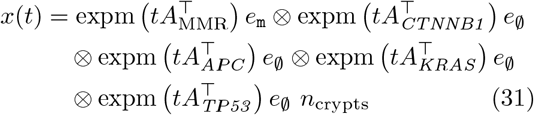

for the case of the first group of patients. This reduces the computational costs tremendously, as only several small matrices have to be considered instead of one large matrix.

## 5 Model Extensions

In the basic model, we assume that all genes are independent of each other. This means in particular that the mutation status of one gene does not affect the status of the other genes. However, mutations change the functional behavior of a cell and thus, specific mutations affect the probability of certain other mutations. In other words, there are mutations which are mutually exclusive or mutations which increase the probability of mutations in other genes [Lei+15].

Instead of changing the adjacency matrix of the basic model, we add the adjacency matrices for the individual extensions to the basic one. In this way, the extended model is a true extension of the basic model. This allows us to study the effects of the different medical processes more precisely.

For the approach presented here, we assume and model the following molecular and biological mechanisms:

**Matrix** *B*: increased point mutation rate of *APC* after MMR deficiency,
**Matrix** *C*: positive association of *CTNNB1* and *MLH1* alterations,
**Matrix** *D*: increased LOH rate after *APC* inactivation,
**Matrix** *E*: mutual enhancement of effects *C* and *D*,
**Matrix** *F*: increased mutation rate of *KRAS* after MMR deficiency.

These matrices are considered in the extended model

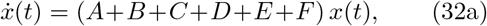

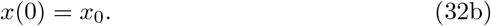

In the following sections, we explain all the extensions in detail.

### 5.1 Increased Point Mutation Rate of *APC* after MMR Deficiency

MMR deficiency leads to an increased mutation rate, especially in microsatellites [Cha03]. Among others, this is true for the point mutation rate of *APC*. Thus, we assume that the point mutation rate of *APC* is increased by a factor *β* + 1 if the crypt has an MMR-deficient state. This is assumed to be independent of the state of the other genes.

As we do not want to change the matrix *A* of the basic model, we introduce an additional matrix *B*. This means, instead of multiplying single entries of *A* by *β* + 1, we add a matrix *B* to *A* with corresponding entries multiplied by *β*.

We define the extension matrix *B* by

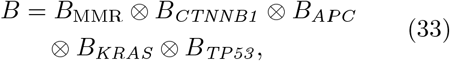

where *B*_*APC*_ is the adjacency matrix of the gene mutation graph in Figure 4 and

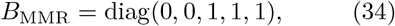

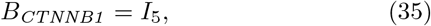

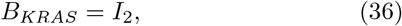

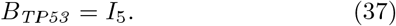

**Figure 4:**
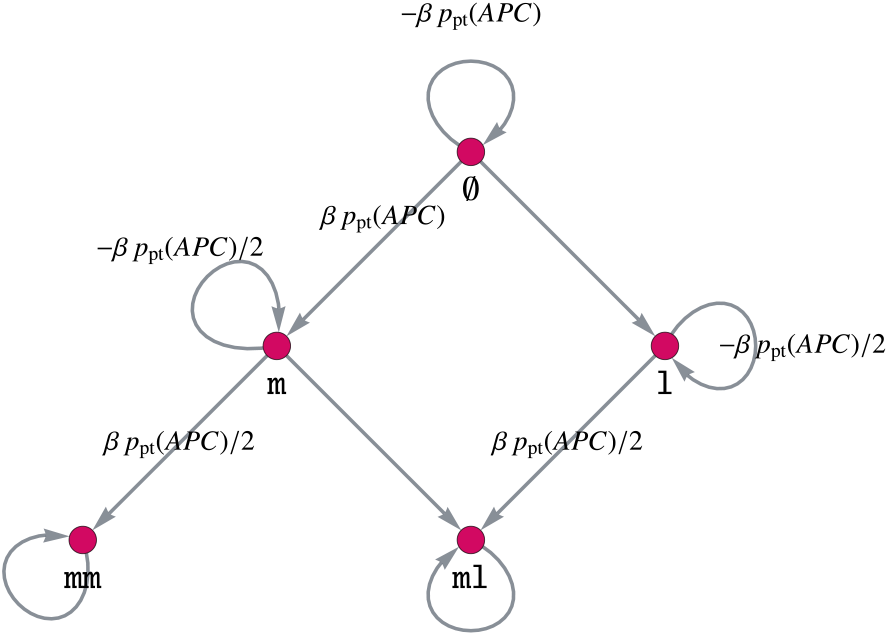
**Gene Mutation Graph of *APC*** for the model extension of increasing the point mutation rate of *APC* after MMR deficiency.

Here, 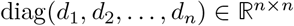 denotes a diagonal matrix with entries *d*_*i*_, *i* ∈ [*n*] on its diagonal.

The definition (33) of the extension matrix *B* yields the desired result of increasing the point mutation rate of *APC* after MMR deficiency. This can be explained intuitively: We only want to increase the point mutation rate after MMR deficiency, meaning that the MMR state should be `mm`, `ml` or `ll`, leading to the matrix *B*_MMR_. Further, this influence of MMR on *APC* is independent of the other genes, meaning that it should hold for all states of the other genes. Thus, we choose the respective identity matrices for *KRAS*, *CTNNB1* and *TP53* and connect all matrices via the Kronecker product, instead of the Kronecker sum in the basic model.

### 5.2 Positive Association of *CTNNB1* and *MLH1* Alterations

According to [Eng+20], somatic *CTNNB1* mutations are significantly higher in *MLH1*-cancers than in the other MMR gene-associated colorectal cancers. For illustration purposes, we assume that this might be due to an inactivation of both genes by an LOH event with an occurrence rate *r*_effLOH_, which we set to *r*_effLOH_ = 0.9. In order to depict this dependency, we introduce an additional matrix *C* which is based on a combined gene mutation graph for *MLH1* and *CTNNB1* and its connection with the remaining genes via the Kronecker product. Note that this is possible due to the chosen ordering of the genes.

The matrix 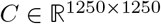 is given by

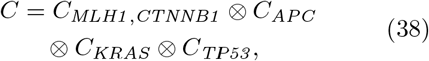

where *C*_*APC*_ = *C*_*TP53*_ = *I*_5_ and *C*_*KRAS*_ = *I*_2_. The matrix *C*_*MLH1,CTNNB1*_ is the adjacency matrix corresponding to the combined gene mutation graph for *MLH1* and *CTNNB1*. We explain in the following how this combined gene mutation graph is built and illustrate it in Figure 5.

**Figure 5:**
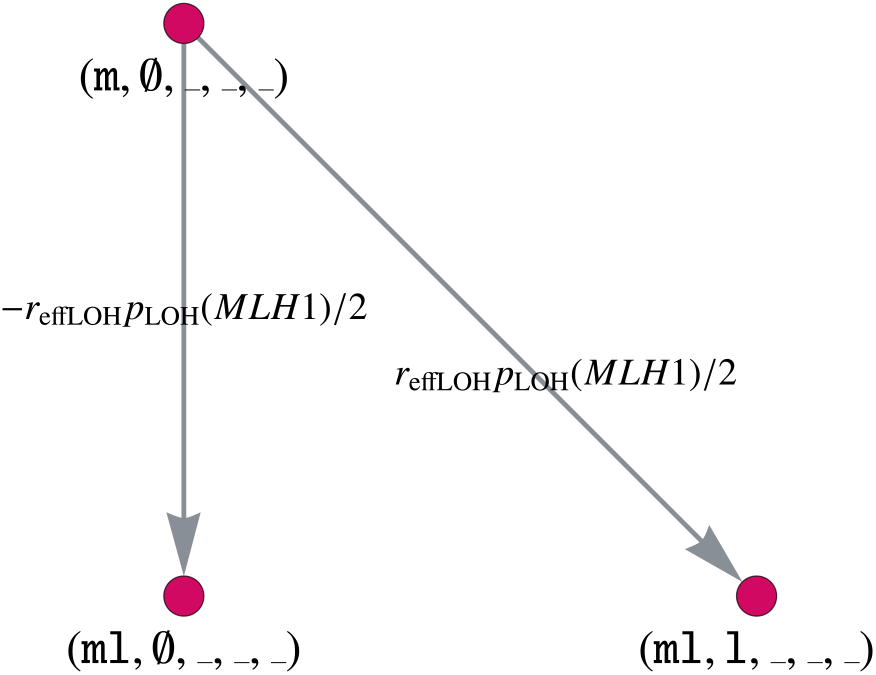
Model Extension For the Positive Association of *MLH1* and *CTNNB1*. Part of the combined gene mutation graph for *CTNNB1* and *MLH1* of the extension *C*. The gene mutation graphs for the other possible gene states *MLH1* ∈ {`l, ll`}, *CTNNB1* ∈ {`m, ml`} are defined in an analogous way.

Let denote an arbitrary state of the corresponding gene. Instead of multiplying the edge weight *p*_LOH_(MMR)/2 of the edge (`m`, ∅, ‒, ‒, ‒) → (`ml`, ∅, ‒, ‒, ‒) by (1 − *r*_effLOH_) in the original matrix *A*, we add a matrix *C* with a corresponding edge weight *r*_effLOH_ *p*_LOH_(MMR)/2. The following edges are added to the matrix *C* with the same weight:

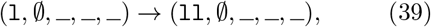

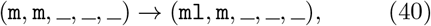

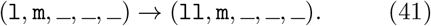

Furthermore, we need to insert the following new edges with edge weight *r*_effLOH_ *p*_LOH_(*MLH1*)/2

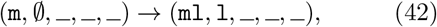

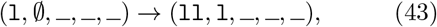

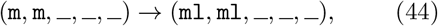

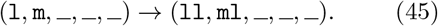

All other entries of *C* are zero, leading to a sparse matrix with only 400 non-zero entries.

### 5.3 Increased LOH Rate after *APC* Inactivation

The following extension of the basic model deals with the increased LOH rate of *APC*-inactivated crypts, which is assumed to be the case in many tumors [Now+02]. In the latter, we will denote those *APC*-inactivated crypts by *APC*−/−, which are inactivated due to `mm` or `ml`.

As further LOH events can occur for MMR, *CTNNB1* and *TP53* in *APC*−/− crypts, we have to introduce individual matrices for each effect leading to the extension matrix *D* = *D*_1_ + *D*_2_ + *D*_3_, where

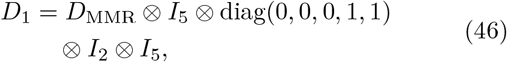

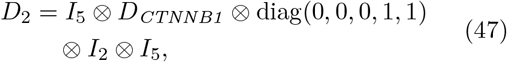

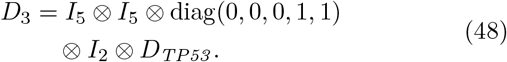

Analogous to the extension *B*, we define a gene mutation graph of MMR, *CTNNB1* and *TP53* with parameter *δ* such that the LOH rate is increased by a factor *δ* + 1. This is illustrated in Figure 6 for *CTNNB1* and *TP53*, where the gene mutation graph for MMR is defined analogously.

**Figure 6:**
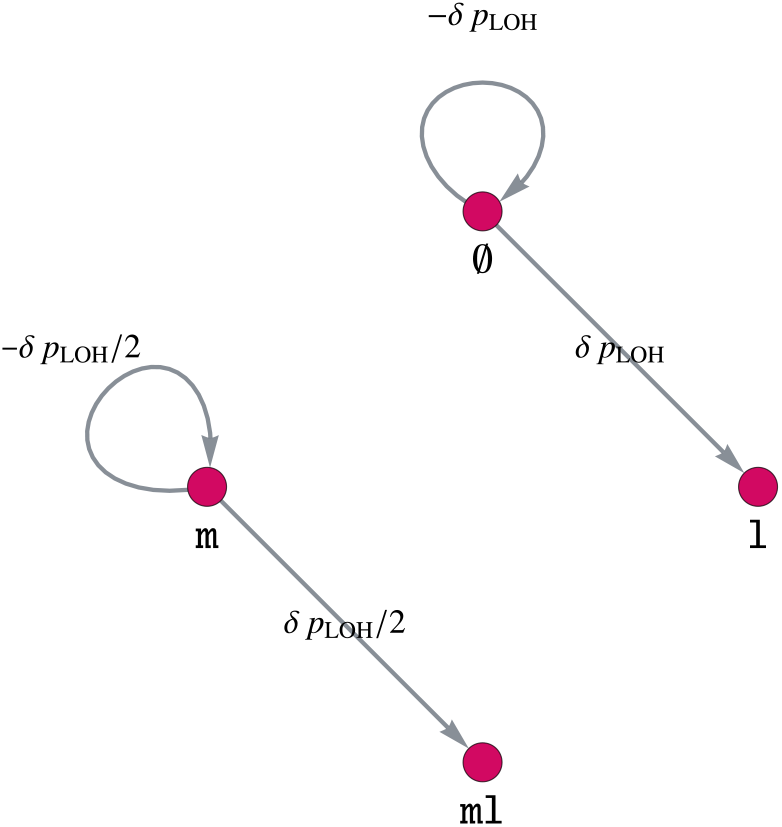
Model Extension for Increasing the LOH Rate of MMR, *CTNNB1* and *TP53* by a Factor *δ* + 1 After *APC* Inactivation. Gene mutation graph for both genes, *CTNNB1* and *TP53*, of the extension *D*. The gene mutation graph for MMR is defined in an analogous way.

### 5.4 Mutual Enhancement of Effects *C* and *D*

*APC* inactivation increases the LOH rate of other genes, including *MLH1*, which is modeled by extension matrix *D*. Further, there is a positive association of *MLH1* and *CTNNB1* alterations, which we can model in the same way as an LOH event, as described in extension matrix *C*. Thus, we would like to demonstrate how to model the mutual enhancement of two effects, which will be described by an additional matrix *E*. As for the matrix *C*, we build the combined adjacency matrix for *MLH1* and *CTNNB1* and combine it with the other genes via the Kronecker product, i.e.

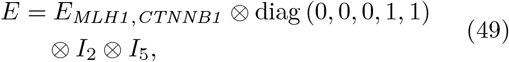

where again, the ordering is essential to enable an efficient implementation.

This enhancement only affects the *APC*−/− crypts, thus we use diag(0, 0, 0, 1, 1) for the *APC* matrix. Analogous to Figure 5, we illustrate parts of the gene mutation graph for the combination of *MLH1* and *CTNNB1* after *APC* inactivation in Figure 7.

**Figure 7:**
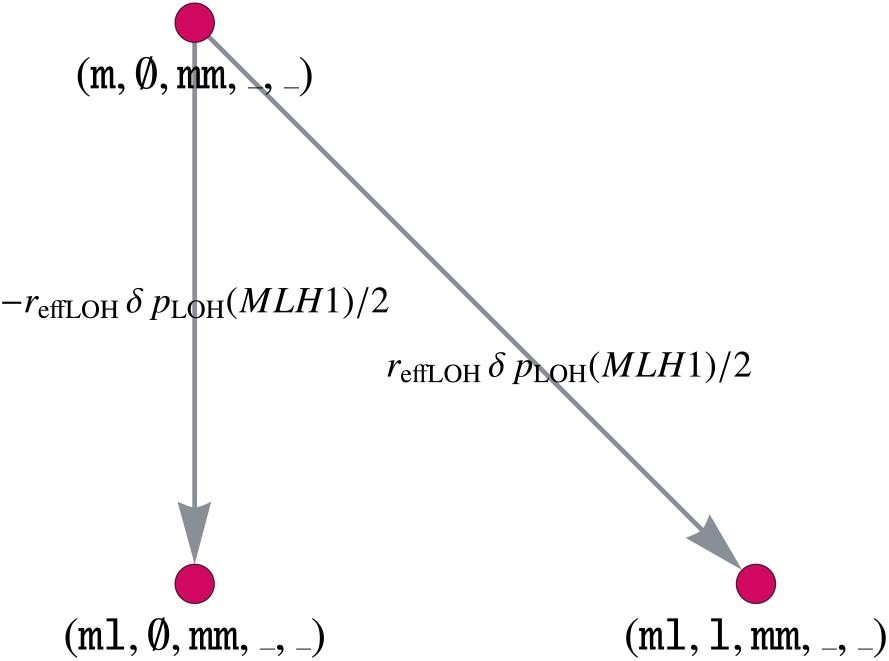
Model Extension for the Mutual Enhancement of Two Extensions by a Factor *δr*_effLOH_. Part of the gene mutation graph for *CTNNB1* and *MLH1* after *APC* inactivation considered by the extension *E*. The gene mutation graphs for the other possible gene states *MLH1* ∈ {`l`, `ll`}, *CTNNB1* ∈ {`m`, `ml`}, *APC* ∈ {`ml`} are defined in an analogous way.

### 5.5 Increased Mutation Rate of *KRAS* After MMR Deficiency

*KRAS* is an oncogene with one point mutation sufficient for activation, where mainly codon 12 or 13 are hit. Codon 13 mutations are known to be associated with and enriched in MMR-deficient cancers, as these mutations are more likely to occur under the influence of MMR deficiency [Aha+18]. We will consider this association by increasing the *KRAS* mutation rate after MMR deficiency by a factor *ζ* + 1. For this, the extension matrix *F* is defined analogously to the extension matrix *B* with the corresponding matrix entries multiplied by *ζ*. The gene mutation graph of *KRAS* is given in Figure 8.

**Figure 8:**
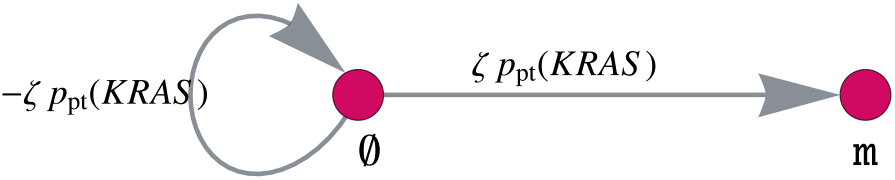
Model Extension for Increasing the Mutation Rate of *KRAS* After MMR Deficiency. Gene mutation graph of *KRAS* for the model extension matrix *F* with the *KRAS* mutation rate increased by a factor *ζ*.

## 6 Modifications to the Model

In Section 4 and 5, we introduced a mathematical modeling approach for colorectal carcinogenesis using the example of Lynch syndrome. We will present modifications to the model to handle other forms of colorectal carcinogenesis such as Lynch-like and MSS carcinogenesis, as well as colorectal carcinogenesis in FAP patients.

For example, this can be done by changing the initial values of the model to differentiate between sporadic and hereditary cases or to consider germline mutations in different genes, e.g. MMR in Lynch syndrome and *APC* in FAP.

Further, we can include other mutation states of already included genes, for instance the wild-type state in the MMR gene for the Lynch-like and sporadic MSI case, and we can adapt specific parameters to account for specific carcinogenesis mechanisms like we will do for the example of FAP later in this section.

Finally, we describe the potential for modifications to account for cancer evolution in other organs.

### 6.1 Non-Lynch and FAP

#### Lynch-like and Lynch Syndrome Carcinogenesis

The main difference between Lynch-like and Lynch syndrome carcinogenesis is the absence or presence of a monoallelic MMR germline variant as a first hit at birth. In Lynch syndrome carcinogenesis, all body cells, including those constituting colonic crypts, already carry a monoallelic mutation in one of the MMR genes, whereas in Lynch-like carcinogenesis all cells start with wild-type MMR genes. By introducing the additional vertex ∅ in 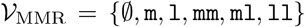 with point mutation and LOH rates described in Sections 4.2 and 4.3, it is possible to represent those two forms of MSI carcinogenesis. The initial value changes by *x*_0_ = 0 except for the entry corresponding to (`m`, ∅, ∅, ∅, ∅) or (`l`, ∅, ∅, ∅, ∅) in the hereditary case and (∅, ∅, ∅, ∅, ∅) in the sporadic case for which the value is set to *n*_crypts_.

#### MSS Carcinogenesis

It is possible to model the evolution of MSS colorectal cancers with the proposed model by not including MMR genes in the vertex set. Due to the absence of MMR in the model, *CTNNB1* mutations should be much less frequent. The classical adenoma-carcinoma model including *APC*, *KRAS* and *TP53* should be the dominant pathway of carcinogenesis.

#### FAP Carcinogenesis

Another application of the model is the evolution of colorectal cancers in a second hereditary syndrome, namely familial adenomatous polyposis (FAP). Those patients have a single germline mutation in *APC*, which is known to be a point mutation in almost all cases [NN93; Ras+16]. Thus, the dynamical system starts with all crypts in the state (∅, ∅, `m`, ∅, ∅).

As reported in [Gry09], we assume that the germline mutations are not equally distributed among the base pairs of the *APC* gene. Instead, they are concentrated at specific codons leading to the fact that we change the number of hotspot base pairs in the FAP case. Due to [KV96], the classical FAP case is associated with germline mutations in codons 1250 − 1464, leading to the assumption *n*_hs_ = 600 in our model for FAP simulations. Thus by changing the parameters of the model, we are able to model other cases of colorectal carcinogenesis.

The common regions of germline mutations described above are also correlated with the most occurring polyps (more than 5,000) [KV96] in FAP patients. With an estimated diameter of 4.8 mm per polyp [Gol+03] and 0.09 mm per crypt [Sta+15], this would result in 10^7^ crypts in a polypous state. Thus, our model simulations should also reflect that the number of polyps, assumed to consist of *APC*−/− crypts, should be much higher than in the sporadic case.

### 6.2 Cancer in Other Organs

In general, it is possible to modify the model in such a way that it can not only model carcinogenesis in the colon but also in other organs. For this, the incorporated genes have to be changed as well as the definitions of point mutations and LOH events have to be adapted to account for different cell structures. The application to other organs will be considered in future work.

## 7 Results

We present the results of modeling the evolution of human colorectal crypts in a typical Lynch syndrome patient over the course of 70 years. The model starts with a germline mutation in MMR in all crypts at birth and yields the temporal evolution of the crypt distribution among all genotypic states, where we only show the results for *MLH1* and *MSH2*, as those are related to the highest colorectal cancer incidence in Lynch syndrome [Eng+20].

### 7.1 Evolution of Crypts with Specific Genotypic States

Making use of equation (17), we extracted and combined different genotypic states from the overall distribution. We did so for MMR-deficient crypts as well as other more advanced states, which we refer to adenomatous and cancerous states. They are defined in the following way:

**MMR-deficient:** MMR-deficient; *CTNNB1*, *APC*, *KRAS*, *TP53* intact, i.e. (`mm`, ∅, ∅, ∅, ∅) + (`ml`, ∅, ∅, ∅, ∅) + (`ll`, ∅, ∅, ∅, ∅)
**State 1:** MMR-proficient or MMR-deficient, *CTNNB1* activated; *APC* inactivated; *KRAS* and *TP53* intact (called early adenomatous)
**State 2:** MMR-proficient or MMR-deficient, *CTNNB1* activated; *APC* inactivated; *KRAS* activated; *TP53* intact (called late adenomatous)
**State 3:** MMR-proficient or MMR-deficient, *CTNNB1* activated; *APC* and *TP53* inactivated; *KRAS* activated (called cancerous)

The parameters are set in such a way that the number of MMR-deficient crypts is quantitatively comparable to the clinical data presented in [Sta+15]. We show the results for *MLH1* and *MSH2* in Figure 9.

**Figure 9:**
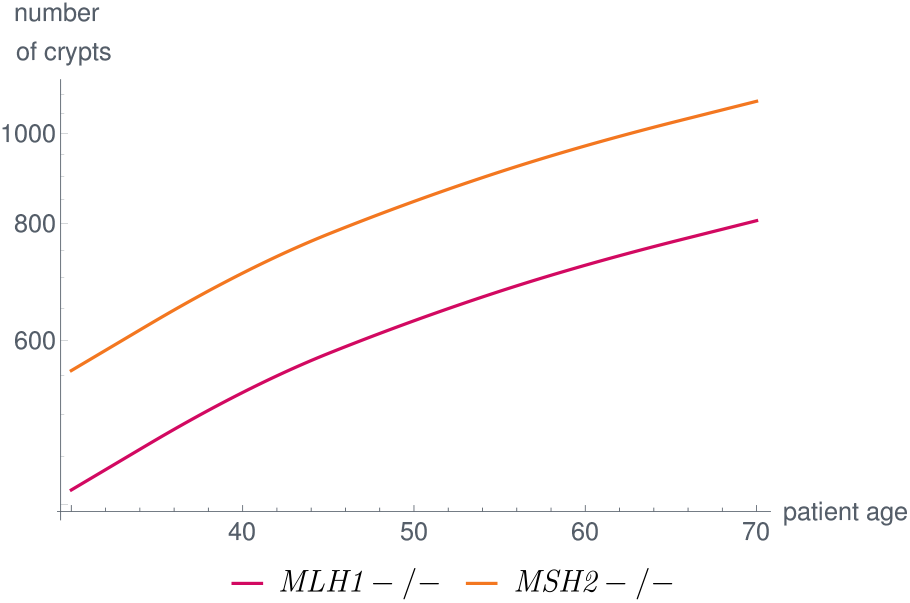
Number of MMR-deficient Crypts Over the Life of a Typical Lynch Syndrome Patient for *MLH1* and *MSH2*. The parameters in the model are set in such a way that the simulation results are in concordance with published data [Sta+15]. In our model, differences among genes are due to differences in coding region and gene lengths as well as the magnitude of the effects of the extended mechanisms.

Further, the results for early and advanced adenomatous and cancerous states are given in Figure 10 for a typical Lynch syndrome patient with a germline mutation in *MLH1*. It is important to note that we can analyze e.g. the relative contribution of MMR-deficient and MMR-proficient adenomatous and cancerous states. With the chosen parameter combinations, this relative contribution changes between the advanced adenomatous and the cancerous states. We will further elaborate these contributions in Section 7.3. Further, it is possible to compare the evolution of these states with respect to the contribution of *APC* and *CTNNB1*. Note that some of the parameters are chosen without any bio-molecular data at hand meaning that some of the absolute numbers of crypts presented here may not match the real numbers if measurable. As soon as further data are available either for the parameters or for the evolution of crypt numbers over time or both, parameter learning will be possible.

**Figure 10:**
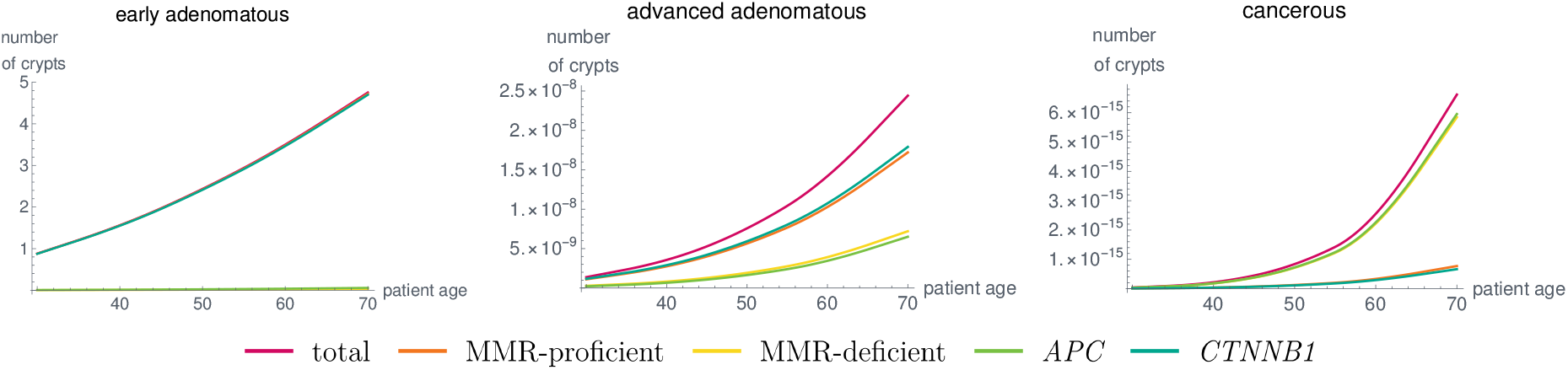
**Number of Crypts Over Time In a Typical *MLH1* Carrier in Combined States**, like early adenomatous, advanced adenomatous and cancerous states as defined in the text for the given parameter set. Due to the extensions of the model accounting for different genetic dependencies, the distribution of MMR-deficient and MMR-proficient, as well as the contribution of *APC* and *CTNNB1* change for the different states. Due to the lack of suitable medical data, parameter learning was not performed in a rigorous way. As soon as data are available, this can be done using different mathematical techniques.

### 7.2 Influences of Variants in MMR Genes

The model is able to compare the carcinogenesis process for the different MMR genes in order to examine gene-specific differences. This in particular includes the questions of whether and how the distribution of crypts in various states changes when considering different MMR genes. More generally, the distribution among the different pathways of Lynch syndrome carcinogenesis may vary among the MMR genes. As the different pathways need different treatment and surveillance strategies, it is essential for Lynch syndrome-related clinical guidelines to examine the gene-specific associations with the pathways, as depicted in [Eng+20].

An early example is given in Figure 9 showing the differences among MMR-deficient crypt foci which are the first detectable precursor lesions of the Lynch syndrome carcinogenesis pathways 2 and 3 illustrated in the Graphical Abstract. Differences among the MMR genes are reported for adenoma and carcinoma incidences of Lynch syndrome patients [Eng+20]. In the model, the differences are due to differences in the properties of the MMR genes, such as coding region and gene lengths, and due to the fact that the extensions of the model influence the evolution of the crypts differently. As soon as there are more data available on bio-molecular mechanisms or there are further pathogenic variant hypotheses to be tested, these differences can be made even more explicit by introducing additional extensions and factors. This will be the subject of future work.

### 7.3 Distribution Among the Carcinogenesis Pathways

We analyzed the proportion of MMR-proficient and MMR-deficient crypts in various states to determine the proportion in which MMR deficiency occurred as an initial event in carcinogenesis of Lynch syndrome carriers. The results are shown in Figure 11 and are similar to the currently available data [Aha+18] with a slight underestimation of MMR-deficient *APC*−/− crypts compared to MMR-proficient ones.

**Figure 11:**
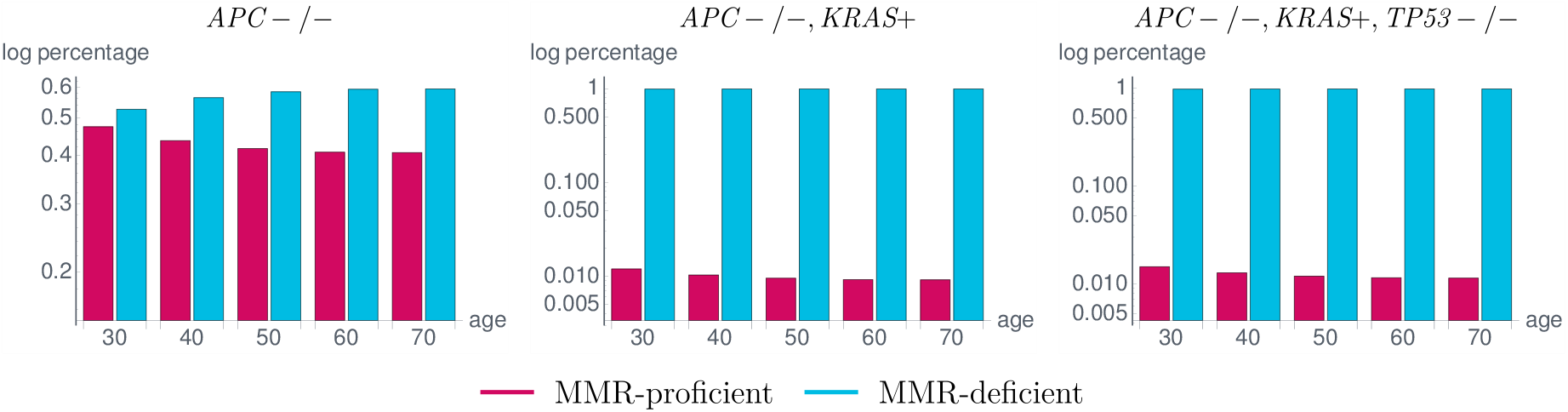
**Proportion of MMR-proficient and MMR-deficient Crypts in a Typical *MLH1* Carrier in Different States** corresponding to the states in the classical adenoma-carcinoma sequence by Vogelstein [VK93]. Among the *APC*−/− crypts (left), the number of MMR-deficient crypts is up to 20% higher than the number of MMR-proficient ones. This difference largely increases with the subsequent *KRAS* activation (*KRAS*+) (middle) and *TP53* inactivation (*TP53*−/−) (right) leading to the fact that almost all crypts in the last state, corresponding to a cancerous state, are MMR-deficient. These simulation results are in concordance with available data with a slight underestimation of MMR-deficient *APC*−/− crypts [Aha+18].

In general, in the basic model, the distributions in Figure 11 are the same as there are no influences between the different genes. In the extended model, we can recognize the dependencies, as the distributions vary within the subsequent states. From *APC*−/− to *APC*−/− and *KRAS*-activated crypts, the difference in the proportions of MMR-proficient and MMR-deficient crypts greatly increases with the given parameter setting leading to the fact that almost all *APC*−/−, *KRAS*-activated crypts are MMR-deficient. As more of the *APC*−/− crypts are MMR-deficient, this seems to imply that MMR deficiency is often the initial event in Lynch syndrome carcinogenesis.

Further, the proportions do not change if *TP53* inactivation happens because currently, there is no such effect incorporated in the extended model for e.g. increasing the mutation rate of *TP53* after MMR deficiency or after *KRAS* activation.

### 7.4 Analysis of Parameter Contributions

The results were obtained with the set of parameters given in Table 3. We analyzed the influences of the parameters on the simulation results. First, the number of point mutations *n*_pt_, the number of cells *n*_cells_, and the number of crypts *n*_crypts_ determine the absolute values of the analyzed numbers.

**Table 3:**
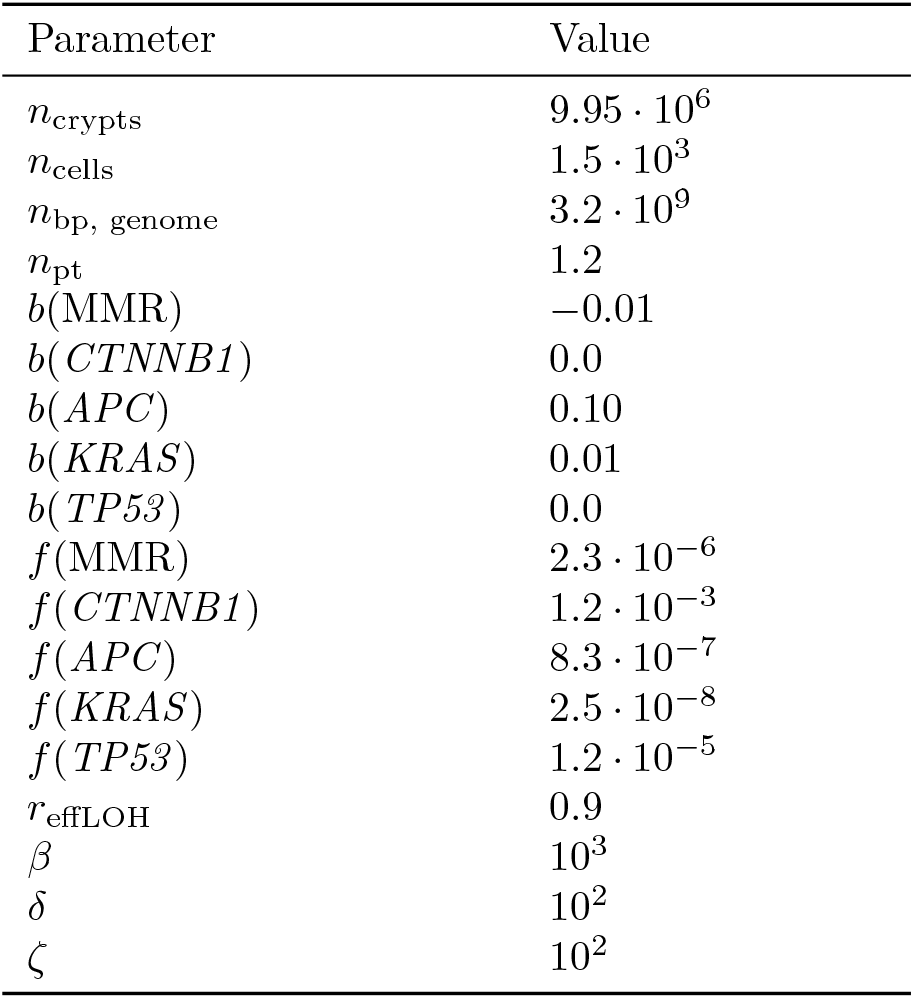
Parameter setting for the shown results.

| Parameter | Value |
| --- | --- |
| $n_{\text{crypts}}$ | $9.95 \cdot 10^6$ |
| $n_{\text{cells}}$ | $1.5 \cdot 10^3$ |
| $n_{\text{bp, genome}}$ | $3.2 \cdot 10^9$ |
| $n_{\text{pt}}$ | 1.2 |
| $b(\text{MMR})$ | -0.01 |
| $b(CTNNB1)$ | 0.0 |
| $b(APC)$ | 0.10 |
| $b(KRAS)$ | 0.01 |
| $b(TP53)$ | 0.0 |
| $f(\text{MMR})$ | $2.3 \cdot 10^{-6}$ |
| $f(CTNNB1)$ | $1.2 \cdot 10^{-3}$ |
| $f(APC)$ | $8.3 \cdot 10^{-7}$ |
| $f(KRAS)$ | $2.5 \cdot 10^{-8}$ |
| $f(TP53)$ | $1.2 \cdot 10^{-5}$ |
| $r_{\text{effLOH}}$ | 0.9 |
| $\beta$ | $10^3$ |
| $\delta$ | $10^2$ |
| $\zeta$ | $10^2$ |

Further, the relation of the hotspot length and the gene length determines the relative frequency of point mutations and LOH events for the individual genes, which can be changed by the model extensions for specific genotypic states. Here, the magnitude of the parameters *r*_effLOH_, *β, δ*, and *ζ* determines how large the contribution of the individual extensions is.

The parameters *b*(gene) affect the slope of the crypt evolution curve. In our case, *b*(MMR) < 0 leads to the fact that further MMR-deficient crypts are disadvantageous for the crypt survival leading to fewer additional MMR-deficient crypts with increasing age (Figure 9).

In contrast, *APC* inactivation is modeled as an advantage for the crypts such that *b*(*APC*) > 0 leads to more additional *APC*-inactivated crypts with increasing age.

Furthermore, the relation of the fixation affinities *f* (gene) for different genes seems to influence the ordering of the mutations. A larger value of *f* (gene) leads to a faster fixation in this gene and thus to an earlier event in carcinogenesis (Figure 11).

As soon as there are more molecular data available, parameter learning could be applied to the model in order to get a deeper understanding of the underlying mechanisms in Lynch syndrome carcinogenesis. In particular, there is still uncertainty in the data about the fitness advantages and disadvantages of individual genetic changes as well as on the fixation affinities of mutations. General information on mutational dependencies and how they affect the phenotype of the cells is crucial to extend the model with further bio-molecular mechanisms.

### 7.5 Non-Lynch and FAP

We compared different types of colorectal carcinogenesis by changing the initial values of the dynamical system or by adapting other parameters.

First, we compared the number of MMR-deficient crypts in Lynch-like and Lynch syndrome patients, as illustrated in Figure 12. The latter is much larger in Lynch syndrome patients than in Lynch-like patients, corresponding with [Sta+15].

**Figure 12:**
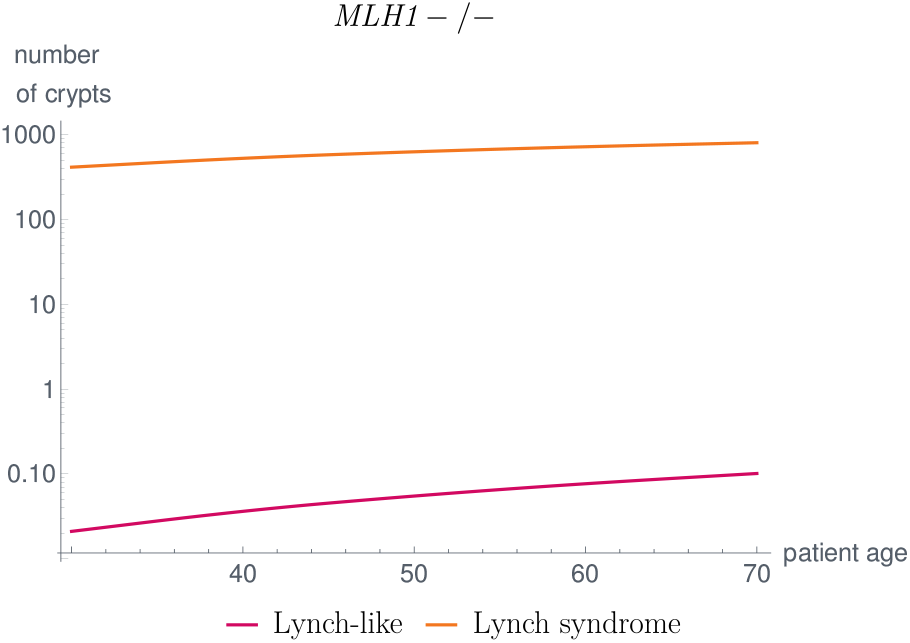
Comparison of MMR-deficient Crypts in Lynch-like and Lynch Syndrome Patients. The number of MMR-deficient crypts is significantly higher in Lynch syndrome patients compared to Lynch-like patients, which matches the findings in [Sta+15].

This is due to the fact that in Lynch syndrome, a germline mutation in one allele of the MMR gene is already present such that an additional mutation leading to MMR-deficiency could be gained earlier in life.

Further, we compared the *APC*−/− crypt evolution of a typical FAP patient with a sporadic case without a germline mutation in *APC* for all crypts. We used the parameter setting given in Table 3, except for *n*_hs_(*APC*) = 600. We changed the number of hotspot base pairs in the FAP case due to the fact that the germline mutations are not equally distributed among the base pairs of the *APC* gene, as described in Section 6.1.

With the given parameter set, our model simulations yield between 10^4^ − 10^5^ *APC*−/− crypts, which is below the estimates calculated from the literature (see Section 6.1). The time evolution of the number of crypts is shown in Figure 13. It would be necessary for the future to obtain age-dependent data as well as further measurements to be able to adapt the parameters accordingly.

**Figure 13:**
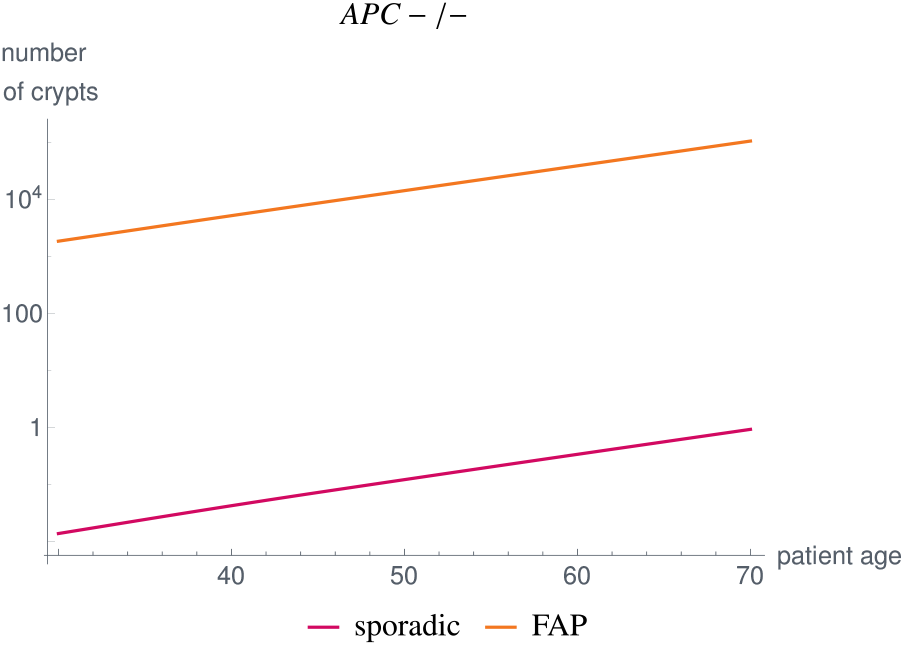
**Comparison of *APC*−/− Crypts In the Sporadic Case and In FAP Patients**, where we changed the initial value of the dynamical system as well as *n*_hs_(*APC*) = 600 for FAP. Our simulation results yield numbers below estimates found in the literature [Gol+03; KV96; Sta+15]. With improved measurements, future work will adapt the parameters accordingly.

## 8 Conclusion and Outlook

We presented a mathematical model for the multiple pathways in colorectal carcinogenesis based on dynamical systems with a Kronecker structure, which models the number of colorectal crypts being present in different genotypic states. We focused on the evolution of key genotypic states occurring in Lynch syndrome, the most common inherited colorectal cancer syndrome, namely alterations in the MMR genes, *CTNNB1*, *APC*, *KRAS* and *TP53*.

With a certain probability, a cell within a crypt is prone to mutations in a specific gene. We assumed that the mutations are spread evenly across the whole genome meaning that the mutation rate of a gene within this cell is proportional to the length of the gene and the total number of mutations occurring in a cell during cell division. As there are multiple cells within a crypt each having an individual cell cycle, it takes some time until the mutation is present in the whole crypt, a process called fixation. Further, a mutation could be washed out of the crypt, if it is not advantageous enough for fixation to occur. Thus, we assumed that the mutation rate of a gene in a crypt also depends on a fixation tendency of the specific genetic event. The edge weights in the graph representation correspond to the mutation rates between those genotypic states of crypts, where the mutation rates are computed based on the described assumptions. Further, we distinguish the genetic alterations between point mutations and loss of heterozygosity events.

The presented modeling approach consists of a basic model and several extensions. The basic model forms the framework of the system assuming that all genetic alterations are independent of each other. There is no gene influencing the genetic state of another gene.

Mathematically, this is represented by building a mutation graph for each individual gene and combining them using the Cartesian graph product. This means that the matrix of the corresponding dynamical system can be obtained by combining the adjacency matrices using the Kronecker sum. As mutations cannot be reverted, the model matrix is an upper triangular matrix leading to even more efficient algorithms for solving the system.

The model extensions represent specific correlations and dependencies of genetic events to account for a more realistic illustration of Lynch syndrome carcinogenesis. They are chosen in concordance with existing medical hypotheses and data, e.g. the increased point mutation rate after MMR deficiency which is one of the reasons for increased lifetime risk of colorectal cancer in Lynch syndrome patients.

The matrices of the model extensions again have a Kronecker structure. Further, they are added to the basic model in order to keep the latter unchanged. This eases the analysis of the individual effects on the overall model solution. If further medical hypotheses and data are available, it is simple to include further extensions in the model.

In addition, the model can be easily modified to other types of carcinogenesis, such as sporadic MMR-deficient cancers, often called Lynch-like cancers, other hereditary colorectal cancers like familial adenomatous polyposis, and microsatellite-stable colorectal cancers.

In principle, it is possible to apply the model structure to other organs by modifying the mutation probability definitions according to the underlying cell structure and by incorporating different genes with appropriate model extensions based on the predominant genetic effects. This will be the subject of further investigation.

Further, we model carcinogenesis on the basis of the number of crypts being present with specific genotypic states. The latter can be combined easily with clinically relevant stages like early adenoma, late adenoma, and carcinoma. An early adenoma is often referred to as *APC*−/− crypts. However, the other genes are often not analyzed appropriately meaning that it is not clear whether single somatic mutations not leading to a phenotypic change are already present. This means that it is not clear which genotypic states of the model have to be taken together for a clinically relevant stage. If more molecular data with the analysis of all possible driver genes are available, a comparison of the model with these data will allow for parameter learning of the as yet unmeasurable parameters.

